# Endothelial expression of human APP leads to cerebral amyloid angiopathy in mice

**DOI:** 10.1101/2020.07.22.215418

**Authors:** Yuriko Tachida, Saori Miura, Rie Imamaki, Naomi Ogasawara, Hiroyuki Takuwa, Naruhiko Sahara, Akihiro Shindo, Yukio Matsuba, Takashi Saito, Naoyuki Taniguchi, Yasushi Kawaguchi, Hidekazu Tomimoto, Takaomi Saido, Shinobu Kitazume

**Affiliations:** Disease Glycomics Team, Glycobiology Research Group, Global Research Cluster, RIKEN, Saitama 351-0198, Japan; Preparing Section for New Faculty of Medical Science, Fukushima Medical University School of Medicine, Fukushima 960-1295, Japan; Department of Infectious Disease Control, International Research Center for Infectious Diseases, The Institute of Medical Science, The University of Tokyo, Tokyo 108-8639; Department of Functional Brain Imaging Research, National Institute of Radiological Sciences, Chiba 263-8555, Japan; Departments of Neurology, Graduate School of Medicine, Mie University, Mie 514-8507, Japan; Laboratory for Proteolytic Neuroscience, RIKEN Brain Science Institute, Saitama 351-0198, Japan; Department of Neurocognitive Science, Institute of Brain Science, Nagoya City University Graduate School of Medical Sciences, Nagoya 467-8601, Japan

## Abstract

The deposition of amyloid β (Aβ) in blood vessels of the brain, known as cerebral amyloid angiopathy (CAA), is observed in more than 90% of Alzheimer’s disease (AD) patients. The presence of such CAA pathology is not as evident, however, in most mouse models of AD, thereby making it difficult to examine the contribution of CAA to the pathogenesis of AD. Since blood levels of soluble amyloid precursor protein (sAPP) in rodents are less than 1% of those in humans, we hypothesized that endothelial APP expression would be markedly lower in rodents, thus providing a reason for the poorly expressed CAA pathology. Here we generated mice that specifically express human APP770 in endothelial cells. These mice exhibited an age-dependent robust deposition of Aβ in brain blood vessels but not in the parenchyma. Crossing these animals with APP knock-in mice led to an expanded CAA pathology as evidenced by increased amounts of amyloid accumulated in the cortical blood vessels. These results show that both neuronal and endothelial APP contribute cooperatively to vascular Aβ deposition, and suggest that this mouse model will be useful for studying disease mechanisms underlying CAA and for developing novel AD therapeutics.

## Introduction

Alzheimer’s disease (AD) is a progressive neurodegenerative disorder in humans and the leading cause of dementia. The pathology of AD is characterized by deposition of amyloid β peptide (Aβ) in the brain parenchyma and blood vessels. Aβ is generated by the two-step proteolytic cleavage of amyloid precursor protein (APP), β-site APP cleaving enzyme-1 (BACE1)^1^, ^2^ and γ-secretase^3^, the last of which is composed of four transmembrane proteins (presenilin, nicastrin, Pen2, and Aph1)^4^. When APP is alternatively cleaved at the α-site within the Aβ sequence, subsequent γ-secretase cleavage cannot produce neurotoxic Aβ peptide. Alternative mRNA splicing generates three major APP isoforms, APP695, APP751, and APP770, each of which exhibits cell-specific expression. In this way, neurons express APP695^3^, while vascular endothelial cells and platelets express APP770^5, 6^, and APP751 shows a relatively ubiquitous expression pattern^7^.

Cerebral amyloid angiopathy (CAA) is defined as the deposition of Aβ to the vascular walls of the meninges and brain, giving rise to intracerebral hemorrhage and cognitive impairment in the elderly^8^. However, epidemiological and clinicopathological data alone do not fully describe the pathological association between CAA and AD.

Impaired clearance and/or overproduction of Aβ leads to its accumulation in the brains of AD patients^9^. Most neuronal Aβ is transported across the blood-brain barrier (BBB) into the blood^10^, and as such the blood Aβ can derive from a mixture of APP sources including from neurons and other tissues, all of which can contribute to vascular Aβ deposition. Accumulating evidence shows that several anti-Aβ immunotherapies reduce the amyloid burden in brain parenchyma, while augmenting that of vascular Aβ accumulation^11, 12^, thereby indicating that Aβ accumulation in the brain parenchyma and vessels is regulated in different ways. A major therapeutic concern with some anti-Aβ antibodies is the possible breakdown of the blood-brain barrier (BBB) associated with brain edema and cerebral microhemorrhages, which are observed under magnetic resonance imaging as amyloid-related imaging abnormalities (ARIAs)^13^. To this end, the manner by which brain Aβ accumulates in the vascular walls is an important issue to be clarified. Unfortunately, most AD model mice with parenchymal Aβ deposition exhibit only moderate Aβ deposition in the brain blood vessels^14^.

Based on a preliminary study showing that the level of serum sAPP in rodents is less than 1% of that compared with human serum, we explored the possibility that limited endothelial APP expression could result in a much-reduced CAA pathology in rodent models. In this study, we have developed a novel CAA mouse model that expresses human APP770 in endothelial cells. We show that the mice exhibit vascular Aβ deposition without parenchymal Aβ plaque formation accompanying breakdown of the BBB with aging.

## Results

### Endothelial APP expression in mice

We first measured blood sAPP levels in rats, and found that plasma and serum sAPP levels were 70 pg/ml and 1.1 ng/ml, respectively (Fig. 1a). These levels are markedly lower (0.3 %) than those measured in human samples, in which plasma and serum sAPP770 levels were 47 and 410 ng/ml, respectively. Serum sAPP770 levels are substantially higher than plasma sAPP770 levels because platelets store sAPP770 in α-granules and release it into the serum upon activation^6, 15^. In mice, blood sAPP levels were even undetectable in some cases. Even though APP is considered to be expressed ubiquitously^16^, we hypothesized that endothelial APP expression might be limited in rodents, and that this scenario could result in markedly lower levels of blood sAPP and poor CAA pathology in most AD mouse models. To test this hypothesis, we developed a new mouse model that specifically expresses human APP770 with the Swedish (KM670/671NL) mutation^17^ in vascular endothelial cells. We first generated floxed hAPP770_NL_ mice (EC-APP770^−^) under the CMV early enhancer/chicken β actin promoter (Fig. 1b, c) and then crossed these mice with *Tie2-Cre* mice, which express Cre recombinase in the endothelial cells by *Tie2* promoter^18^, to obtain double Tg mice (EC-APP770^+^). Immunohistochemical analyses of brain sections from these mice showed that the human APP770 signal is found in endothelial marker PECAM positive-cells in EC-APP770^+^ mice but not in EC-APP770^−^ mice (Fig. 1d). We next used brains from EC-APP770^+^ and EC-APP770^−^ mice to isolate endothelial cells, these being positive for PECAM and negative for the neuronal marker β III-tubulin (Fig. 1e). Western blot analysis showed that an hAPP770 signal was not detectable in brain lysates from either type of mouse. In contrast, we could detect an hAPP770 signal in the lysates of brain endothelial cells isolated from EC-APP770^+^ mice but not from EC-APP770^−^ mice. Taken together, these data indicate that hAPP770 is specifically expressed in the endothelial cells of EC-APP770^+^ mice. Using an anti-APP antibody that detects the carboxy-terminal region of APP that is common to both mouse and human APP, we estimated the expression level of endothelial hAPP770 in EC-APP770^+^ mice to be about three times higher than that in EC-APP770^−^ mice.

**Figure 1.**
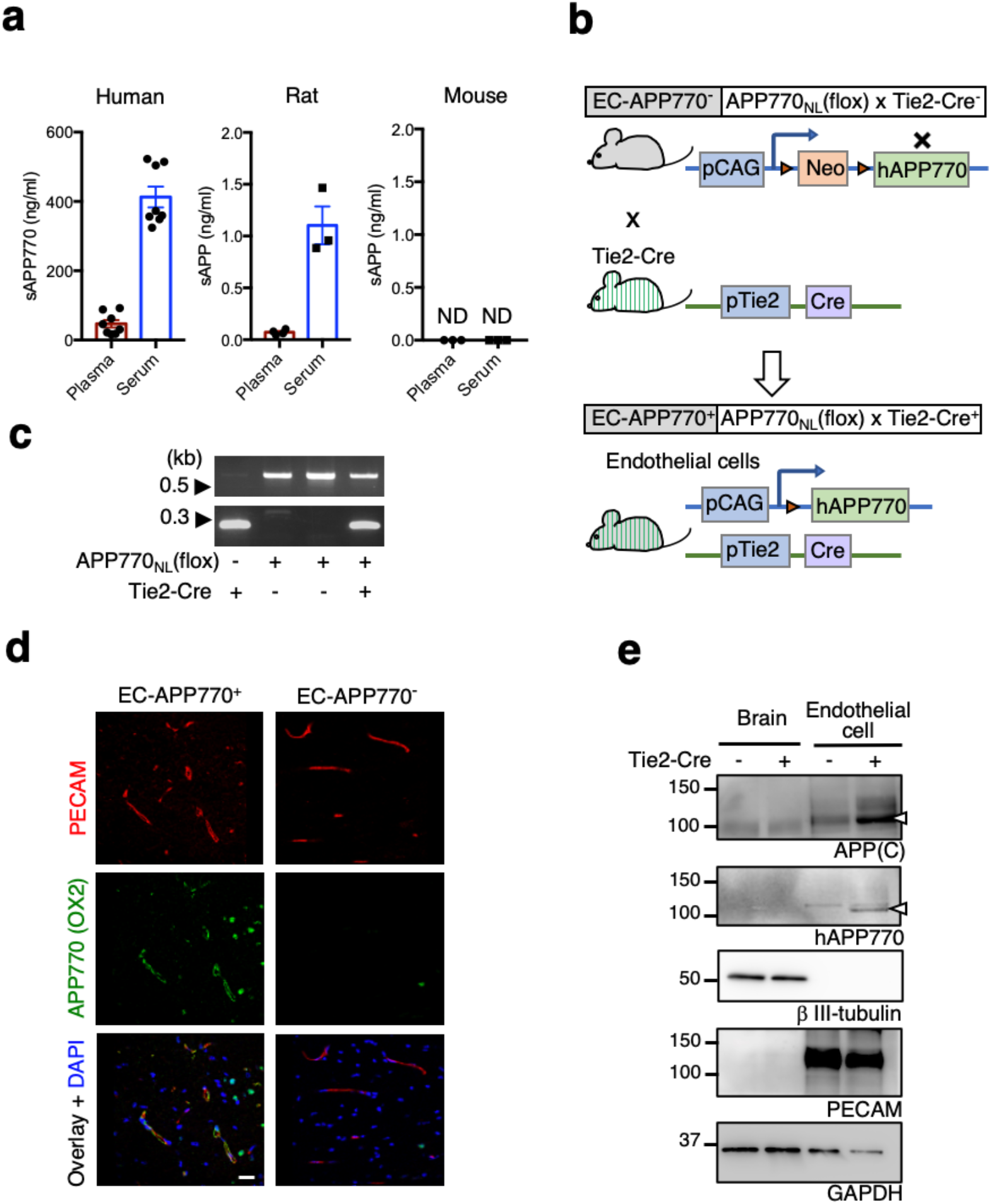
Endothelial APP expression mice. (a) Biochemical measurement of blood sAPP levels in human, rat, and mouse samples. Plasma and serum levels of sAPP770 in humans (n =8) and sAPP in rats (n=3) and mice (n=3) were respectively quantified by sandwich ELISA. Data show mean±s.e.m. (b) Schematic drawing showing endothelial-specific hAPP770 expression in mice. (c) Mouse genotype identified by PCR. In the top image, the 650 bp band is derived from the APP770 floxed allele. The bottom image shows the Tie2-Cre transgene (230 bp band). (d) Endothelial human APP770 expression in EC-APP770^+^ mice. Immunohistochemical analyses of cortical sections of EC-APP770^+^ and EC-APP770^−^ mice with anti-hAPP770 and anti-PECAM antibodies. Scale bar: 20 μm. (e) Biochemical examination of hAPP770 expression in endothelial cells. Lysates from the brains or isolated endothelial cells from EC-APP770^+^ and EC-APP770^−^ mice were analyzed by western blotting for APP, hAPP770, βIII-tubulin, PECAM and GAPDH.

### Aged EC-APP770^+^ mice show Aβ deposition in brain blood vessels

Since a considerable amount of the extracellular region of endothelial APP770 is cleaved for secretion^5^, human sAPP770 should be detectable in the serum of EC-APP770^+^ mice. Indeed, serum sAPP770 levels in these mice were ~7.6 μg/ml whereas they were undetectable in EC-APP770^−^ mice (Fig. 2a). We also measured sAPP770β levels in EC-APP770^+^ mice to be ~90 ng/ml, which is less than 2% of total sAPP770. This compares favorably with the case of human plasma, in which the sAPPα level is about 15-fold that of the sAPPβ level^19^. Cultured endothelial cells secrete both Aβ40 and Aβ42 in the culture medium^5^. In EC-APP770^+^ mice, serum Aβ40 was detectable whereas the Aβ42 level was below the detection limit for our analysis. To our knowledge, these data clearly show for the first time that endothelial APP contributes to blood Aβ levels. Two-year-old EC-APP770^+^ mice showed somewhat reduced serum Aβ40 levels as compared with 1-year-old mice.

**Figure 2.**
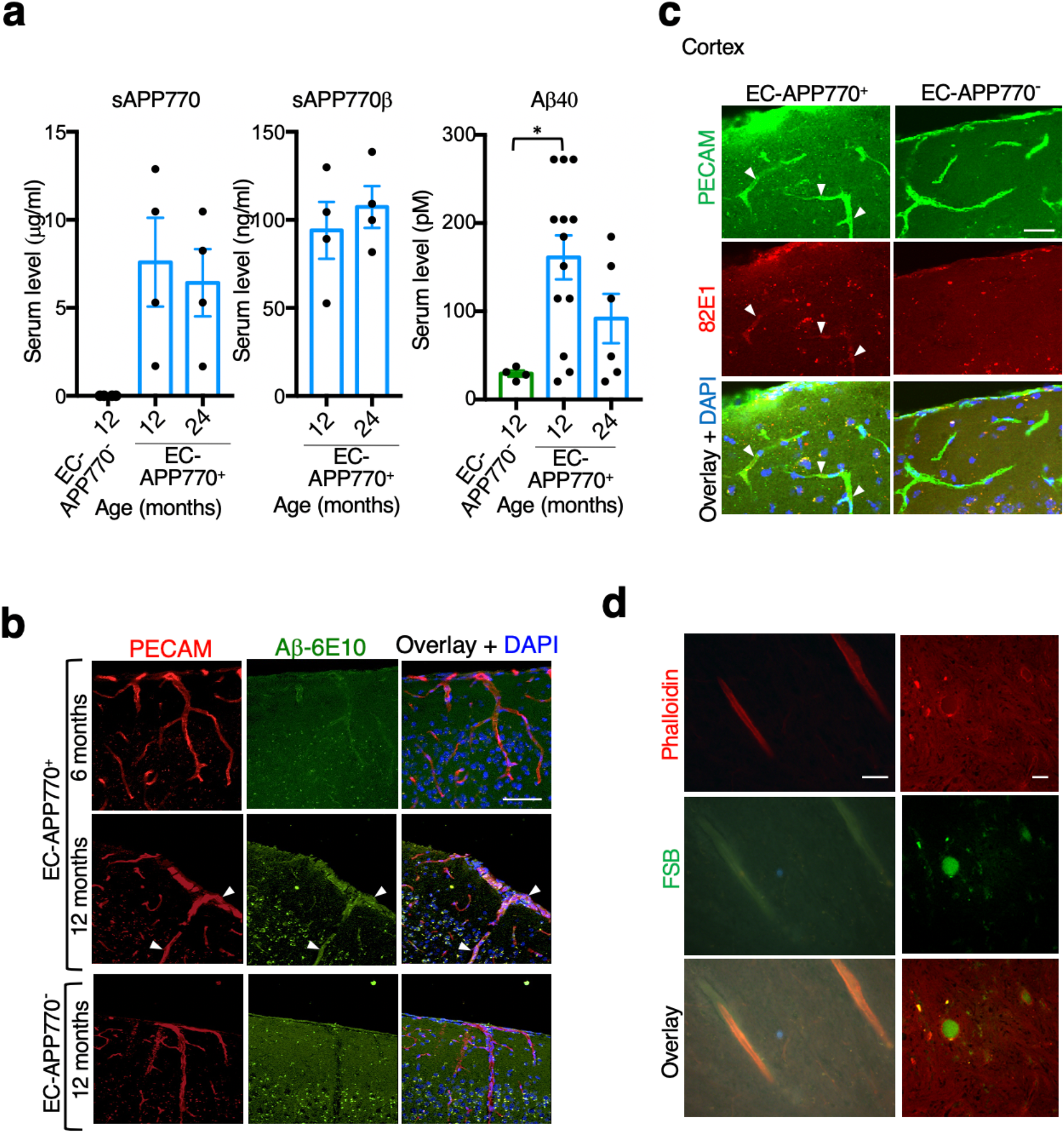
Aged EC-APP^+^ mice show Aβ deposition in brain blood vessels. (a) Serum levels of sAPP770, sAPP770β and Aβ40 in EC-APP770^+^ and EC-APP770^−^ mice are shown as the mean±s.e.m. (n=4-13). (b) Immunohistochemical analysis of brain cortical sections from the EC-APP770^+^ (6 and 12 months old) and EC-APP770^−^ mice (12 months old) using anti-PECAM and anti-Aβ (6E10) antibodies. Arrowheads indicate 6E10-positive endothelial cells. Scale bar: 50 μm. (c) Brain cortex sections from 12-month-old EC-APP770^+^ and EC-APP770^−^ mice were stained by 82E1 for Aβ oligomers and anti-PECAM antibodies. Arrowheads indicate 82E1-positive endothelial cells. Scale bar: 20 μm. (d) Amyloid deposition in cortical blood vessels of 12-month-old EC-APP770^+^ mice was detected with FSB and Alexa546-phalloidin. Scale bar: 20 μm.

We next sought to determine whether EC-APP770^+^ mice show increased CAA pathology with aging. Immunohistochemical analyses showed that EC-APP770^+^ mice exhibit Aβ deposition in the walls of leptomeningeal and cortical arteries at 12 months of age but not at 6 months (Fig. 2b). In contrast, such amyloidosis was absent in EC-APP770^−^ mice even at 12 months. We also employed 82E1 antibody to preferentially detect the Aβ peptides but not full length APP; in this way, 82E1-positive signals were found in the cerebral vessels of EC-APP770^+^ mice but not of EC-APP770^−^ mice at 15 months of age (Fig. 2c). Granular 82E1-positive signals observed in both types of aged mice are considered to be derived from lipofuscin. Furthermore, tail vein injection of 82E1 antibody ^20^ resulted in longer retention of the antibody in the cerebral blood vessels of 22-month-old EC-APP770^+^ mice, compared to EC-APP770^−^ mice (Supplementary Fig. 1). To verify the presence of amyloid plaque deposition in EC-APP770^+^ mice, we used 1-fluoro-2,5-bis(3-carboxy-4-hydroxystyryl)benzene (FSB), which preferentially binds amyloid plaque^21^. In brain sections from 15-month-old EC-APP770^+^ mice, FSB signals were co-localized with the signal for phalloidin-Alexa546, a vascular smooth muscle cell marker (Fig. 2d)^22^. Magnified radial images of blood vessels showed that FSB signals were present in the luminal regions of phalloidin-Alexa546-positive vessels. These observations show that endothelial APP770 expression in mice leads to cerebrovascular Aβ deposits in an age-dependent manner, as we had hypothesized, and that EC-APP770^+^ mice could serve as useful models for studying CAA pathology.

### Deposition of activated complement components in brain blood vessels along with BBB leakage are prominent features in aged EC-APP770^+^ mice

Since the complement system represents a key role for neuroinflammation, and given that complement proteins are associated with cerebral microvascular Aβ deposition^23 24, 25^, we next investigated whether activated complement proteins are localized to cortical blood vessels. In 15-month-old EC-APP770^+^ mice, relatively strong C3b signals were co-localized with PECAM-positive cortical blood vessels (Fig. 3a). C3b binding can activate the membrane attack component (MAC) involving C5b-9. Indeed, C5b-9 signals were significantly higher in the cortical blood vessels of 15-month-old EC-APP770^+^ mice (Fig. 3a, b). C5 convertase containing C3b generates C5a anaphylatoxin. We found that serum C5a levels in EC-APP770^+^ mice were significantly higher than those in EC-APP770^−^ mice (Fig. 3c). Since a BBB leakage is closely associated with CAA ^26^, we next characterized the integrity of the BBB by measuring leakage of fibrin, an endogenous plasma protein ^10^. Perivascular fibrin leakage was observed in a limited number of 12-month-old EC-APP770^+^ mice, whereas none of the aged EC-APP770^−^ mice analyzed showed such fibrin leakage (Fig. 3d). To assess the BBB breakdown in a more quantitative manner, we employed an *in vivo* assay to test blood vessel permeability in brains^27^. Following the intraperitoneal injection of Evans blue, a BBB-impermeant dye that binds to albumin, significant levels of Evans blue were detected in around half of the 15-month-old EC-APP770^+^ mouse brains analyzed (Fig. 3e), but were absent in the brains of similarly aged EC-APP770^−^ mice.

**Figure 3.**
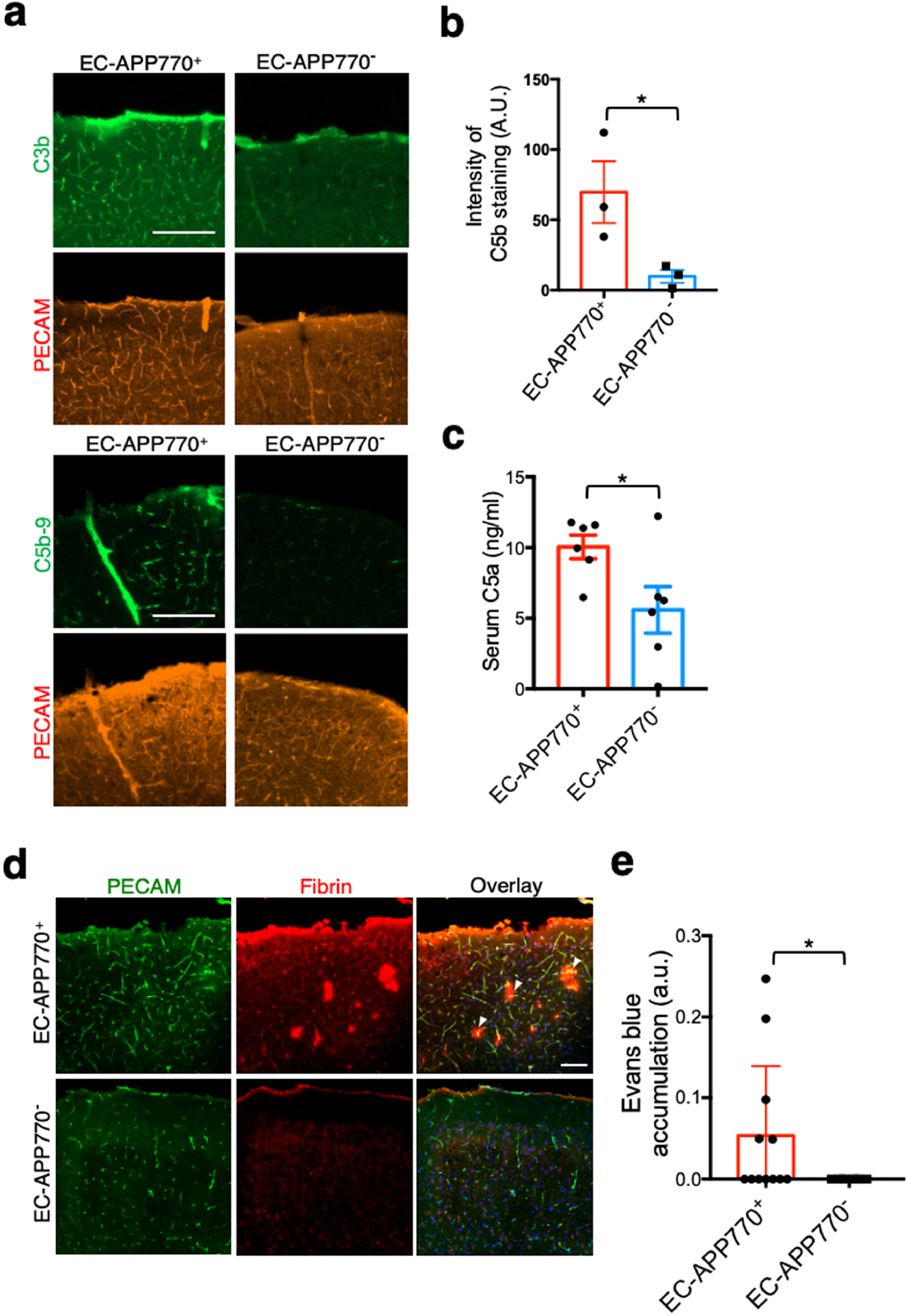
Deposition of activated complement components in brain blood vessels and presence of microhemorrhage are prominent features of aged EC-APP770^+^ mice. (a)Brain cortical sections from 12-month-old EC-APP770^+^ and EC-APP770^−^ mice were stained for C3b and C5b-9 complex with PECAM. Scale bar: 200 μm. (b) Signal intensities of C5b-9 were shown as the mean±s.e.m. (n=3). Student’s *t*-test, p<0.05. (c) Serum C5b levels in 12-month-old EC-APP770^+^ and EC-APP770^−^ mice, determine d by sandwich ELISA, are shown as the mean±s.e.m. (n=6). Student’s *t*-test, p<0.05. (d) Brain cortical sections from 13-month-old EC-APP770^+^ and EC-APP770^−^ mice were stained for fibrin and PECAM. Arrowheads indicate fibrin leakage. Scale bar: 200 μm. (e) Quantification of Evans blue intensity in brains from 13-month-old EC-APP770^+^ and EC-APP770^−^ mice are shown as the mean±s.e.m. (n=8). Student’s *t*-test, p<0.05.

### EC-APP770^+^ mice crossed with APP-KI mice exhibited marked Aβ deposition in blood vessels

We next considered how endothelial APP expression influences AD pathology. We crossed EC-APP770^+^ mice with standard *App*^NL-F/NL-F^ AD model mice and evaluated amyloid plaque formation in FSB-stained brain sections from 15-month-old EC-APP770^+^, EC-APP770^+^:*App*^NL-F/NL-F^, EC-APP770^−^:*App*^NL-F/NL-F^, and *App*^NL-F/NL-F^ mice. In low-magnification images, signals of vascular amyloid deposition were relatively weak in EC-APP770^+^ mice (Fig. 4a). In contrast, *App*^NL-F/NL-F^ mice exhibited weak vascular amyloid deposition in addition to parenchymal amyloid plaque.

**Figure 4.**
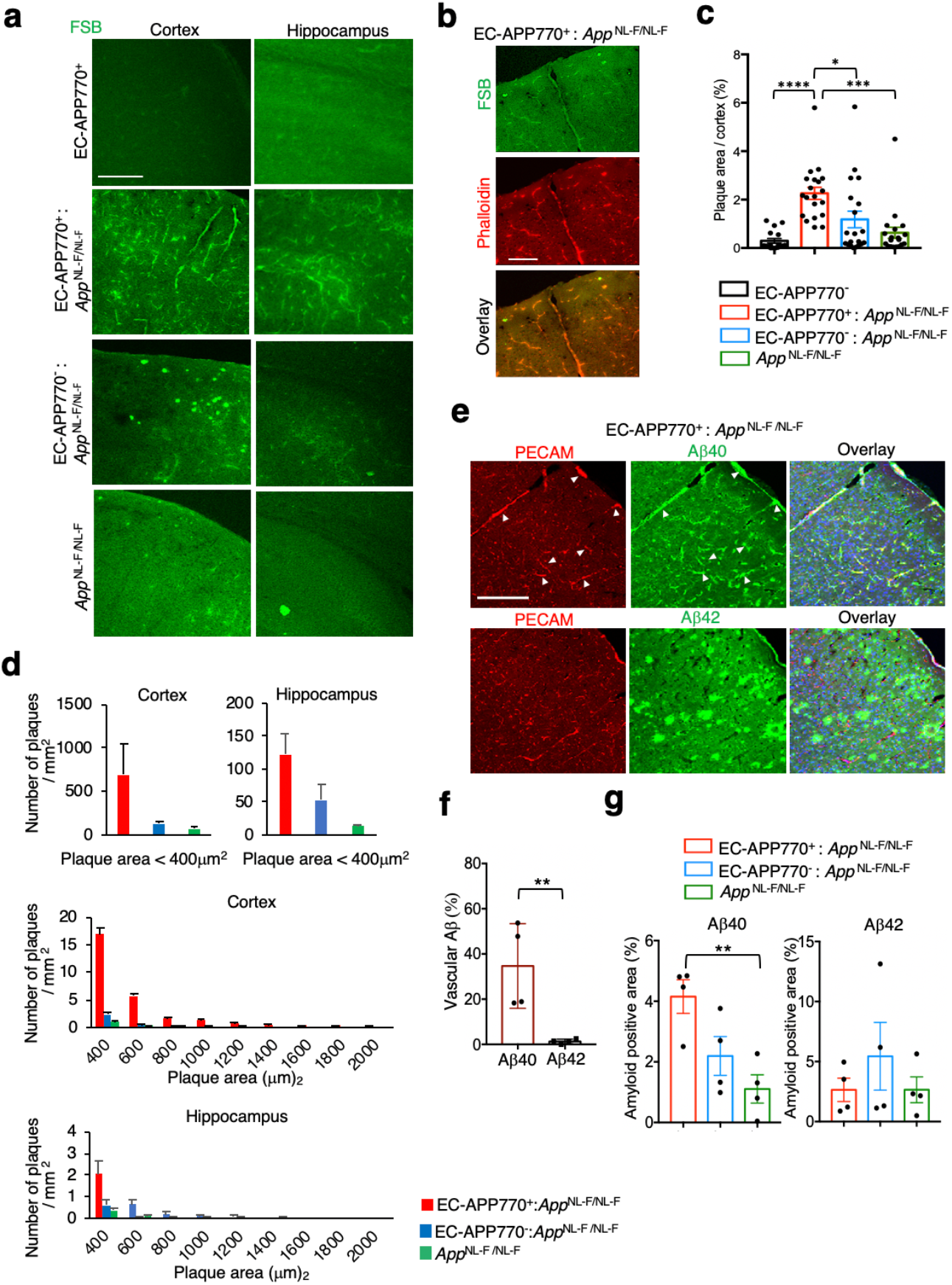
Offspring of EC-APP770^+^ mice crossed with APP-KI mice exhibit marked Aβ deposition in brain blood vessels. Cortical and hippocampal sections from 15-month-old EC-APP770^+^, (a) EC-APP770^+^: *APP*^NL-F/NL-F^, EC-APP770^−^: *APP*^NL-F/NL-F^ and *APP*^NL-F/NL-F^ mice were stained with FSB. Scale bar: 200 μm. (b) Cortical sections from 15-month-old EC-APP770^+^: *APP*^NL-F/NL-F^ mice were stained with FSB and Alexa546-phalloidin. Scale bar: 100 μm. (c) Percentage of FSB-positive plaque area per brain cortex is shown as the mean±s.e.m. (n=4). One-way ANOVA with Tukey’s multiple comparison test, *, p < 0.05; **, p < 0.001. (d) FSB-positive amyloid plaques in cortical and hippocampal sections from 15-month-old EC-APP770^+^ : *APP*^NL-F/NL-F^, EC-APP770^−^ : *APP*^NL-F/NL-F^ and *APP*^NL-F/NL-F^ mice were quantified and classified by size, and are shown in histogram form. (e) Brain cortical sections from 15-month-old EC-APP770^+^ : *APP*^NL-F/NL-F^ mice were stained for Aβ40 and Aβ42 with PECAM. Scale bar: 200 μm. (f) By the analysis of immunostained pictures of EC-APP770^+^: *APP*^NL-F/NL-F^ brain sections, percentage of PECAM-positive Aβ signals relative to total Aβ signals is shown as the mean±s.e.m. (n=4). Student’s *t*-test, p<0.001. (g) Percentage of amyloid positive area per brain cortex is shown as the mean±s.e.m. (n=4). One-way ANOVA with Tukey’s multiple comparison test, **, p < 0.001.

The number of parenchymal plaques in cortical regions of EC-APP770^+^:*App*^NL-F/NL-F^ mice was about two times higher than that in *App*^NL-F/NL-F^ mice (Supplementary Fig. 2). However, as a similar increase was also observed in EC-APP770^−^:*App*^NL-F/NL-F^ mice, we speculated that this could be due to an undesired effect arising from insertion of the floxed hAPP770 transgene. We then decided to determine the integration site of the transgene by next-generation sequencing, and found that it was within the intron between exons 3 and 4 of the *Ikzf3* gene in chromosome 11 (Supplementary Fig. 3a). The integration site was confirmed by genomic PCR using primers against genomic DNA and the transgene, respectively (Supplementary Fig. 3b). The *Ikzf3* gene encodes Aiolos, a member of the Ikaros family of zinc-finger proteins, which shows restricted expression in the lymphoid system, and regulates lymphocyte differentiation^28^. We measured Aiolos expression levels in the spleens of EC-APP770^−^ and wild type mice and found that expression in EC-APP770^−^ mice was 1.4 times higher than that of wild type mice (Supplementary Fig. 3c and d), which could result in a phenotypic alteration in these animals after crossing with AD model mice.

A most striking feature observed in EC-APP770^+^:*App*^NL-F/NL-F^ mice was the massive amyloid plaque deposition in cortical blood vessels (Fig. 4b). The percentage of FSB-positive area in the cortex was highest in EC-APP770^+^:*App*^NL-F/NL-F^ mice (Fig. 4c). In order to characterize amyloid plaque characteristics in greater detail, we performed integrated morphometry analysis in which FSB-positive amyloid plaques were counted and classified on the basis of size. Amyloid deposition to larger vessels, corresponding to plaque sizes over 400 μm^2^, was prominent in EC-APP770^+^:*App*^NL-F/NL-F^ mice (Fig. 4d). As biochemical analyses previously showed that Aβ isoforms shorter than 42 residues form a component of the cerebrovascular amyloid from AD patients^29^, we next performed immunohistochemical analyses on brain sections from EC-APP770^+^:*App*^NL-F/NL-F^ mice to detect different Aβ species. An Aβ40 signal was co-localized with that of PECAM, and thus restricted to brain blood vessels, while an Aβ42 signal was observed exclusively in the brain parenchyma (Fig. 4e, f). Anaysis of EC-APP770^+^:*App*^NL-F/NL-F^, EC-APP770^−^:*App*^NL-F/NL-F^, and *App*^NL-F/NL-F^ mice shows that significantly higher levels of Aβ40 were accumulated in EC-APP770^+^:*App*^NL-F/NL-F^ mice, as compared with *App*^NL-F/NL-F^ mice, while accumulation of Aβ42 is not different in these mice (Supplementary Fig. 4, Fig. 4g).

Taken together, our newly developed mouse model could serve as a useful research tool to study the disease mechanism underlying CAA pathology.

## Discussion

It remains unclear whether the Aβ in accumulated plaque in the cortical blood vessels is derived from the circulation, the blood vessel wall itself, or from the central nervous system ^30^. High levels of plasma Aβ do not cause CAA in transgenic mice expressing signal peptide plus the carboxyl-terminal region of APP that contains Aβ^31^. Nevertheless, as intraperitoneal inoculation with Aβ-rich brain extracts (possibly containing Aβ oligomers) from AD model mice can induce CAA in APP-overexpressing mice^32^, circulating Aβ oligomers may play a seeding or transmission function in vascular Aβ deposition. Neuronal APP-expressing mice also exhibit mild CAA pathology, supporting the notion that neuronal Aβ could be a source of CAA. In this study, we show for the first time that endothelial expression of human APP contributes to vascular Aβ deposition in mice. Remarkably, by crossing with hAPP knock-in mice, in which neuronal APP is expressed, the CAA pathology is markedly exacerbated, also supporting the idea that impaired perivascular drainage of Aβ in the interstitial fluid (ISF) could contribute to CAA pathology^33, 34^. These results indicate that both endothelial APP and neuronal APP co-operatively contribute to the progression of CAA pathology. Several kinds of anti-Aβ antibodies have been developed as AD immunotherapeutics and are currently in the clinical trials^35^. However, administration of anti-Aβ antibodies often causes amyloid-related imaging abnormalities (ARIA), suggestive signals for vasogenic oedema and sulcal effusions (ARIA-E) and microhemorrhage and haemosiderin deposits (ARIA-H), as adverse effects. Since the aged EC-APP770^+^ mice characteristically exhibit vascular abnormalities, such as vascular deposition of activated complement components and enhanced BBB leakage, the mice would be useful to study the degree of ARIA induced by administration of anti-Aβ antibodies and basic molecular mechanism of ARIA.

A disadvantage associated with using EC-APP770^+^ mice is the increased expression of Aiolos due to insertion of the floxed APP transgene within the intron of the gene encoding for Aiolos, *Ikzf3*. Aiolos overexpression has been reported in B cells from patients with systemic lupus erythematosus (SLE)^36^, while it was recently shown that the cerebral small blood vessel disease burden is increased in SLE ^37^. By crossing with AD model mice, both EC-APP770^+^ and EC-APP770^−^ mice exhibited amyloid plaques in the cerebral small blood vessels, an effect that is likely related to the SLE-like features. Further to this, the endothelial-specific angiopoietin 2– Tie2 ligand–receptor system has been identified as a major regulator of endothelial inflammation in SLE. Mating female EC-APP770^−^ mice with wild type male mice should be avoided, since this results in offspring with APP770 expression throughout the body, possibly due to aberrant Tie2 activation in the pregnant mice^38^.

Even though Aβ42 has a higher propensity than Aβ40 to aggregate, immunohistochemical analyses of EC-APP770^+^:*App*^NL-F/NL-F^ mice showed that vascular amyloid consists mostly of Aβ40, as has been reported in the case of AD patients^29^, while parenchymal amyloid consists mostly of Aβ42. One interesting possibility is that endothelial cells in the aged mice could induce specific molecule(s) with high affinity for the Aβ40 oligomer. Brain sections from aged EC-APP770^+^ mice stained with FSB and Alexa546-phalloidin to detect amyloid and vascular smooth muscle cells, respectively, showed that amyloid plaques are mainly located in the luminal side of vascular endothelium. However, longitudinal images of vessel sections showed a vascular FSB signal outside of the smooth muscle cells, possibly due to the disruption of vascular smooth muscle cells, as has been reported in APP overexpressing mice^39^. Indeed, breakdown of the BBB is clearly evident in aged EC-APP770^+^ mice.

Taken together, our newly developed EC-APP770^+^ mice could serve as a useful research tool to study the disease mechanism underlying CAA pathology and the mechanistic interactions between CAA and AD.

## Methods

### Materials

Sources of materials used in this study were as follows: tissue culture medium and reagents, including Dulbecco’s modified Eagle’s medium (DMEM), were from Invitrogen; all other chemicals were from Sigma or Wako Chemicals. Anti-hAPP monoclonal antibody (rat, 1D1)^40^ was conjugated with Alexa 555 using a HyLyte Fluor™ 555 Labeling Kit-NH_2_ (Dojindo). Other commercially available antibodies used were anti-APP (mouse, 22C11, Chemicon), anti-sAPPα (mouse, 6E10, BioLegend), anti-human APP770 (rabbit, Japan-IBL), anti-EEA1 and anti-adaptin γ (mouse, BD Transduction Laboratories), Alexa Fluor 488-anti-Aβ (6E10, BioLegend), anti-PECAM (rat, MEC13.3, BioLegend), anti-tubulin β3 (mouse, TUJ1, BioLegend), anti-CD146 (rat, ME-9F1, BioLegend), anti-GAPDH (mouse, MAB374, Chemicon), anti-FLAG (M2) (mouse, Sigma), and anti-APP (APP(C)), anti-APP OX2, and anti-human sAPPβ (rabbit, IBL-Japan). pCALNL5 (RDB01862, RIKEN BioResource Center) plasmid was from Dr. I. Saito (University of Tokyo).

### Animal Use and Care

All animal experiments were performed at RIKEN, Fukushima Medical University, or the National Institute of Radiological Sciences in compliance with their respective guidelines for animal experiments.

### Generation of EC-APP770^+^ mice

Endothelial APP770-overexpressing mice (EC-APP770^+^) were generated with the help of the Support Unit for Animal Resources Development in RIKEN BSI. The plasmid pCANLN5-hAPP770 was generated by inserting the human APP770 Swedish mutant (hAPP770_NL_) into the pCALNL5 vector^41^. The resulting plasmid has the neomycin-resistant (neoR) gene flanked by two loxP sites under the CMV early enhancer/chicken β actin promoter (CAGp), followed by the hAPP770Sw sequence. pCANLN5-hAPP770 was linearized and injected into C57BL/6J blastocysts. We extracted DNA from ear punches of mouse pups and identified nine F1 generation mice by PCR using the following cocktail of primers^42^: 5’-ATCTCGGAAGTGAAGATG-3’, 5’- ATCTCGGAAGTGAATCTA-3’, 5’-TGTAGATGAGAACTTAAC-3’ and 5’- CGTATAATGTATGCTATACGAAG-3’. Heterozygous EC-APP770^−^ mice were then crossed with Tie2-Cre mice^18^. The *APP770_NL_* allele was identified by PCR (forward, 5’- AGCCGCAGCCATTGCCTTTTATGGTA-3’, reverse, 5’- CAGGTCGGTCTTGACAAAAAGAACCG-3’ for *APP770_NL_*). The *Tie2-Cre* allele was identified by PCR (forward, 5’-ACATGTTCAGGGATCGCCAG-3’, reverse, 5’- TAACCAGTGAAACAGCATTGC-3’). To obtain EC-APP770^+^ and EC-APP770^−^ mice, male EC-APP770^+^ mice were mated with wild type C57BL/6 female mice. EC-APP770^+^ mice were crossed with AD model (*App*^NL-F/NL-F^) mice^42^.

### Determination of the integration site of the APP transgene

The location of the hAPP770 transgene insertion was determined using Illumina NGS (Hokkaido System Science CO., Ltd). The source code to perform the integration site analysis^43^ is available at https://github.com/hbc/li_hiv. Genomic PCR analyses were performed to confirm the insertion loci using F2 (5’- CTTCTGTTGTTTGGCTTTAT-3’), 2A(5’- GACGTCAATGGAAAGTCCCT-3’), R2 (5’-GTTGGTCTCATGAGGCAAGT-3’), 2B (5’- GACGTCAATGGAAAGTCCCT-3’) and R3 (5’-TGTGTGTGGCAAACACTTTG-3’) primers.

### Immunohistochemical and histochemical studies

To prepare brain sections, mice were anesthetized with medetomidine–midazolam–butorphanol and then transcardially perfused with PBS first and with 0.1 M phosphate-buffered 4% paraformaldehyde. Brains were removed and sequentially immersed in the same fixative for 16 h and then phosphate-buffered 30% sucrose for 16 h at 4°C. Sagittal or coronal (30-μm thickness) sections were immunostained using the floating method. Briefly, sections were permeabilized with 0.3% Triton X-100 in 1% BSA-PBS for 30 min first and with 5% goat serum-PBS for 60 min. The sections were then incubated with primary and secondary antibodies and DAPI. Detailed descriptions of the primary and secondary antibodies used are provided in Supplementary Table 1. For the detection of amyloidosis, 1-fluoro-2,5-bis(3-carboxy-4-hydroxystyryl)benzene (FSB) was used^21^. Stained sections were observed using an Olympus FV-1000 confocal microscope. We quantified immunoreactive areas using MetaMorph imaging software (Universal Imaging Corp., West Chester, USA) as previously described^42^. To reduce the variance among tissue sections, data were averaged from at least four sections per mouse.

### *In Vivo* Fluorescence Imaging

Anti-amyloid β (Aβ) oligomer antibody (82E1, IBL) was labeled with Alexa Fluor 555 (Life Technologies) according to the manufacturer’s protocol. Briefly, 100 μl of antibody (1 mg/ml) coupled with Alexa Fluor dye was mixed and incubated in a 100 mM sodium bicarbonate solution for one hour at 25°C. The labeled 82E1 antibody was separated though a purification column. The purified antibody was kept in darkened vials maintained at 4°C before use.

EC-APP770^+^ and EC-APP770^−^ mice (22 months old) were used in this experiment. A chronic cranial window was surgically attached to the bone of the skull as previously described^44^. *In vivo* fluorescence imaging was conducted by two-photon microscopy (TCS-SP5 MP, Leica Microsystems GmbH, Wetzlar, Germany) with an excitation wavelength of 900 nm. The emission signal was separated by a beam splitter (560/10 nm) and detected with a band-pass filter for labeled 82E1 (610/75 nm). A single image plane consisted of 1024 × 1024 pixels, while the in-plane pixel size was 0.25–0.45 μm depending on the instrument zoom factor used. Two-photon imaging was performed prior to and at 1 minute, 1 day and 2 days after the intravenous injection of labeled 82E1 antibody. Volume images were acquired with a z-step size of 2.5 μm up to a maximum depth of 0.2 mm from the cortical surface.

### Isolation of endothelial cells

Endothelial cells were isolated from mouse brains as previously reported^5, 45^, except for the use of anti-CD146 antibody (BioLegend, clone:ME-9F1) coupled with Dynabeads Sheep anti-Rat IgG (Thermo Fisher Scientific).

### Western blot experiments

Cells from mouse brain or spleen tissues were lysed with T-PER Tissue Protein Extraction Reagent (Thermo) containing Complete Protease Inhibitor Cocktail (Roche Applied Science). Protein concentrations were determined with BCA protein assay reagents (Pierce). Samples (25 – 40 μg of protein) were subjected to SDS-PAGE (5–20% gradient gel) and transferred to nitrocellulose membranes. For western blot analyses, following incubation with 5% non-fat dried milk in TBS-containing 0.1% Tween-20, the membranes were incubated with primary antibodies and horseradish peroxidase (HRP)-labeled secondary antibodies. Signals were detected with SuperSignal West Dura Extended Duration Substrate (Thermo Fisher Scientific) using an ImageQuant LAS-4000 Mini imager (GE Healthcare). Intensities of the resultant protein bands were quantified using ImageQuant TL software (GE Healthcare).

### Quantification of sAPP, Aβ

To measure sAPP770, sAPP770β, Aβ40 and Aβ42 in plasma or serum, a Human sAPP Total Assay Kit, an sAPPβ-w Assay Kit (highly sensitive) (IBL-Japan), a Human / Rat βamyloid (40) ELISA kit and a Human / Rat βamyloid (42) assay kit High-Sensitive (Wako) were used, respectively. To measure rat sAPP, a Mouse / Rat sAPPα (highly sensitive) Assay Kit (IBL-Japan) was used. A mouse complement component C5a DuoSet ELISA (DY2150, R&D systems) was used to measure serum C5a in mice.

### Evans blue assay

To assess BBB integrity, mice were administered 0.2 ml of 2 % Evans blue (Sigma) by intraperitoneal injection^46^. Six hours later, the mice were anesthetized with medetomidine– midazolam–butorphanol and perfused with PBS. Brain and lung tissue (100 mg each) homogenates were incubated with 1 ml formamide (Sigma) at 55 °C for 24 h to extract the Evans blue. The formamide / Evans blue mixture was centrifuged, and the supernatant absorbance was measured at 610 nm with a microplate reader.

### Statistical analysis

All data are shown as mean ± SEM. For comparison between two groups, statistical analyses were performed by Student’s *t*-test. Multiple comparison tests were performed using GraphPad Prism 7 software. Normality was tested using a D’Agostino-Pearson normality test. For comparisons among three or more groups, one-way analysis of variance (ANOVA) followed by a post-hoc test (Tukey’s multiple comparison test) or Bonferroni’s multiple comparison test were performed. The data were collected and processed in a randomized and blinded manner.

## Data availability

All relevant data are available from the corresponding authors upon reasonable request. The source data underlying Figs. 1a, 1c, 1e, 2a, 3a, 3d, 4b, 4c, 4g–i, 5b, 5c, and Supplementary figure 2 are provided in the Source data file. Antibodies used for western blot (WB) and immunohistochemical (IHC) analyses are shown in Supplementary Table 1.

## Acknowledgment

We thank M. Yanagisawa (University of Texas Southwestern Medical Center) for providing Tie2-Cre transgenic mice, I. Saito (The Institute of Medical Science, The University of Tokyo) for the pCALNL5 vector, S. Lichtenthaler (DZNE) for anti-hAPP(1D1) antibody, and staff at the BSI Research Resource Center for producing APP770 (flox) mice. This work was supported by AMED (grant number JP18am0101036) to SK, the Mitsubishi Foundation (to SK), and KAKENHI (16K08601 to SK and 25840043 to YT).

## Author contributions

Y.T., S.M., R.I., N.O., H.T., A.S. and Y.M. designed and performed the experiments. S.K. conceived the study, designed the experiments and wrote the manuscript. H.T., T. Saito., Y.K., N.S., N.T., and T. Saido designed the experiments and contributed to data interpretation.

---

**Supplementary Table 1.** Antibodies used in this study.

| **Primary antibody** | **Secondary antibody** | **Application** |
| --- | --- | --- |
| Rabbit anti-APP770<br>(IBL-Japan, #18961, 1:50) | Alexa 488-donkey anti-rabbit<br>(Thermo, A-11008, 1:300 ) | Immunofluorescent detection in<br>Figure 1d |
| Rat anti-mPECAM<br>(Biolegend, MEC13.1, 1:50) | Alexa488-donkey anti-rat<br>(Thermo, A-11006, 1:300 ) | Immunofluorescent detection in<br>Figure 2c, 3c |
| Rabbit anti-APP©<br>(IBL-Japan, #18961, 1:100) | HRP-conjugated anti-rabbit<br>(GE Healthcare, NA934, 1:3000) | Western blot analysis in Figure<br>1e |
| Rabbit anti-hAPP770<br>(IBL-Japan, #18961, 1:100) | HRP-conjugated anti-rat IgG<br>(GE Healthcare, NA935, 1:3000) | Western blot analysis in Figure<br>1e |
| Rabbit anti-beta III tubulin<br>(Abcam, ab18207, 1:400) | HRP-conjugated anti-rabbit<br>(GE Healthcare, NA934, 1:3000) | Western blot analysis in Figure<br>1e |
| Rat anti-mPECAM<br>(Biolegend, MEC13.1, 1:1000) | HRP-conjugated anti-rat<br>(GE Healthcare, NA935, 1:3000) | Western blot analysis in Figure<br>1e |
| mouse anti-GAPDH<br>(Merck, mAB374, 1:1000) | HRP-conjugated anti-mouse<br>(GE Healthcare, NA931, 1:3000) | Western blot analysis in Figure<br>1e |
| Rat anti-mPECAM<br>(Biolegend, MEC13.1, 1:50) | Alexa546-donkey anti-rat<br>(Thermo, A-11030, 1:300 ) | Immunofluorescent detection in<br>Figure 1d, 2b, 3a, 4e |
| Mouse anti-amyloid $\beta$<br>(Biolegend, 6E10, 1:100) | Alexa488-donkey anti-mouse<br>(Thermo, A-11001, 1:300 ) | Immunofluorescent detection in<br>Figure 2b |
| Mouse anti-amyloid $\beta$<br>(IBL-Japan, 82E1, 1:50) | Alexa546-donkey anti-mouse<br>(Thermo, A-11030, 1:300 ) | Immunofluorescent detection in<br>Figure 2c |
| Mouse anti-C3/C3b<br>(Abcam, ab11871, 1:50) | Alexa488-donkey anti-mouse<br>(Thermo, A-11001, 1:100 ) | Immunofluorescent detection in<br>Figure 3a |
| Mouse anti-C5b-9<br>(Abcam, ab66768, 1:100) | Alexa488-donkey anti-mouse<br>(Thermo, A-11001, 1:100 ) | Immunofluorescent detection in<br>Figure 3a |
| Mouse anti-fibrin α-chain IgM<br>(Abcam, ab19079, 1:50) | Alexa546-anti mouse IgM<br>(Thermo A-21045, 1:100) | Immunofluorescent detection in<br>Figure 3c |
| Mouse anti-amyloid $\beta$(1-40)<br>(IBL-Japan, #10047, 1:50) | Biotin-goat anti-mouse IgG<br>(Thermo, 31800, 1:100 ) | Immunofluorescent detection<br>using Tyramide Signal<br>Amplification (T20932) in<br>Figure 4e, S4 |
| Rabbit anti-amyloid $\beta$(1-42)<br>(IBL-Japan, #18582, 1:50) | Biotin-goat anti-rabbit IgG<br>(Thermo, 31820, 1:300 ) | Immunofluorescent detection<br>using Tyramide Signal<br>Amplification (T20932) in<br>Figure 4e, S4 |
| Mouse anti-aiolos<br>(Biolegend, clone 882, 1:1000) | HRP-conjugated anti-mouse<br>(GE Healthcare, NA931, 1:3000) | Western blot analysis in Figure<br>S3c |
| Mouse anti-actin<br>(Sigma, AC-40, 1:1000) | HRP-conjugated anti-mouse<br>(GE Healthcare, NA931, 1:3000) | Western blot analysis in Figure<br>S3c |
| Alexa555-conjugated anti-hAPP<br>(82E1) |  | *In vivo* fluorescence imaging in<br>Figure S1 |

**Supplementary Figure 1.**
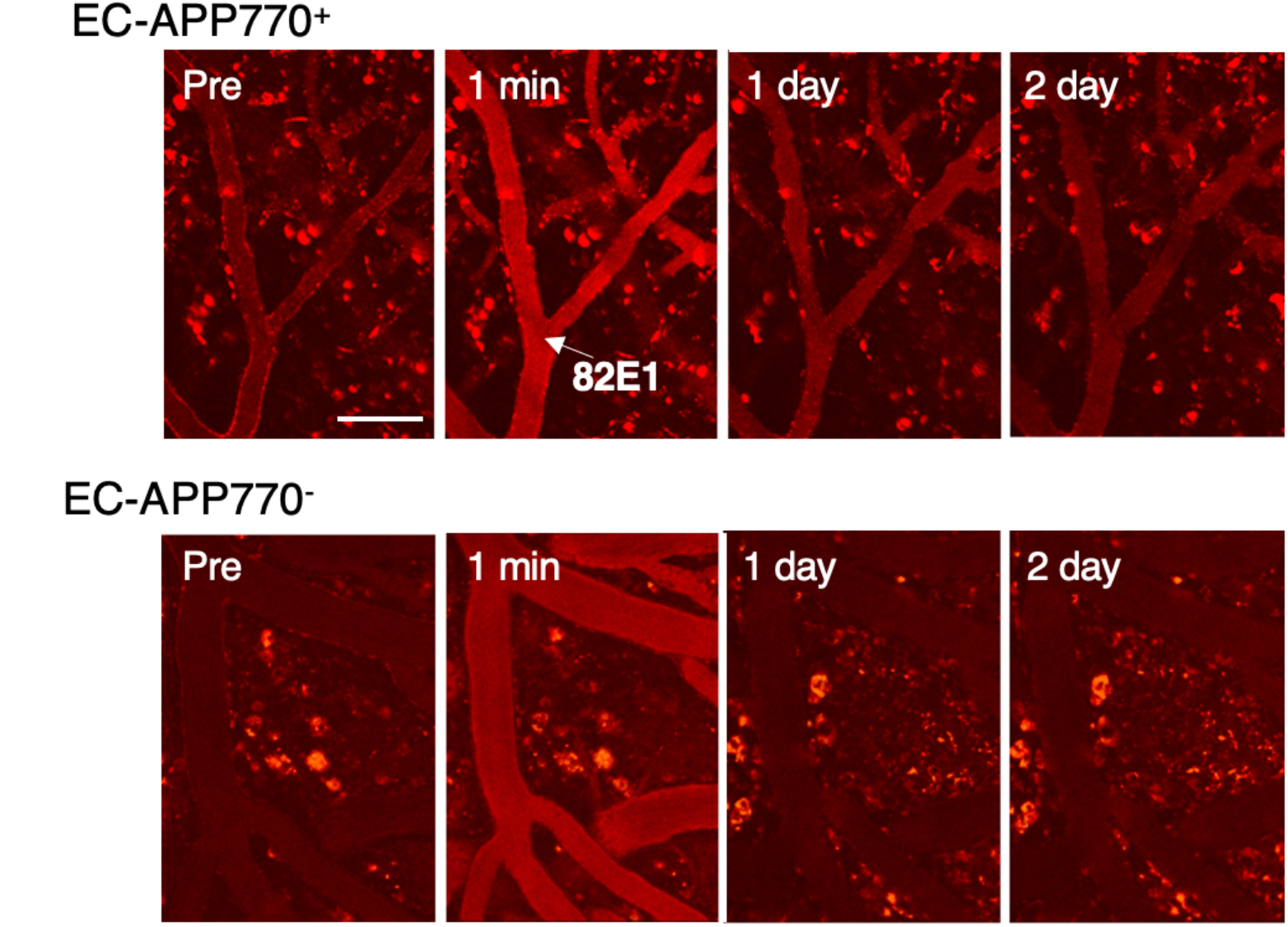
Detection of Aβ oligomer in EC-APP770^+^ mice. To detect cerebrovascular Aβ, *in vivo* fluorescence imaging was performed prior to (Pre) and at 1 minute (1 min), and 1 day (D1) and 2 days (D2) after intravenous injection of labeled 82E1 antibody using 22-month-old EC-APP770^+^ (upper panels) and EC-APP770^−^ (lower panels) mice. Scale bar: 100 μm.

**Supplementary Figure 2.**
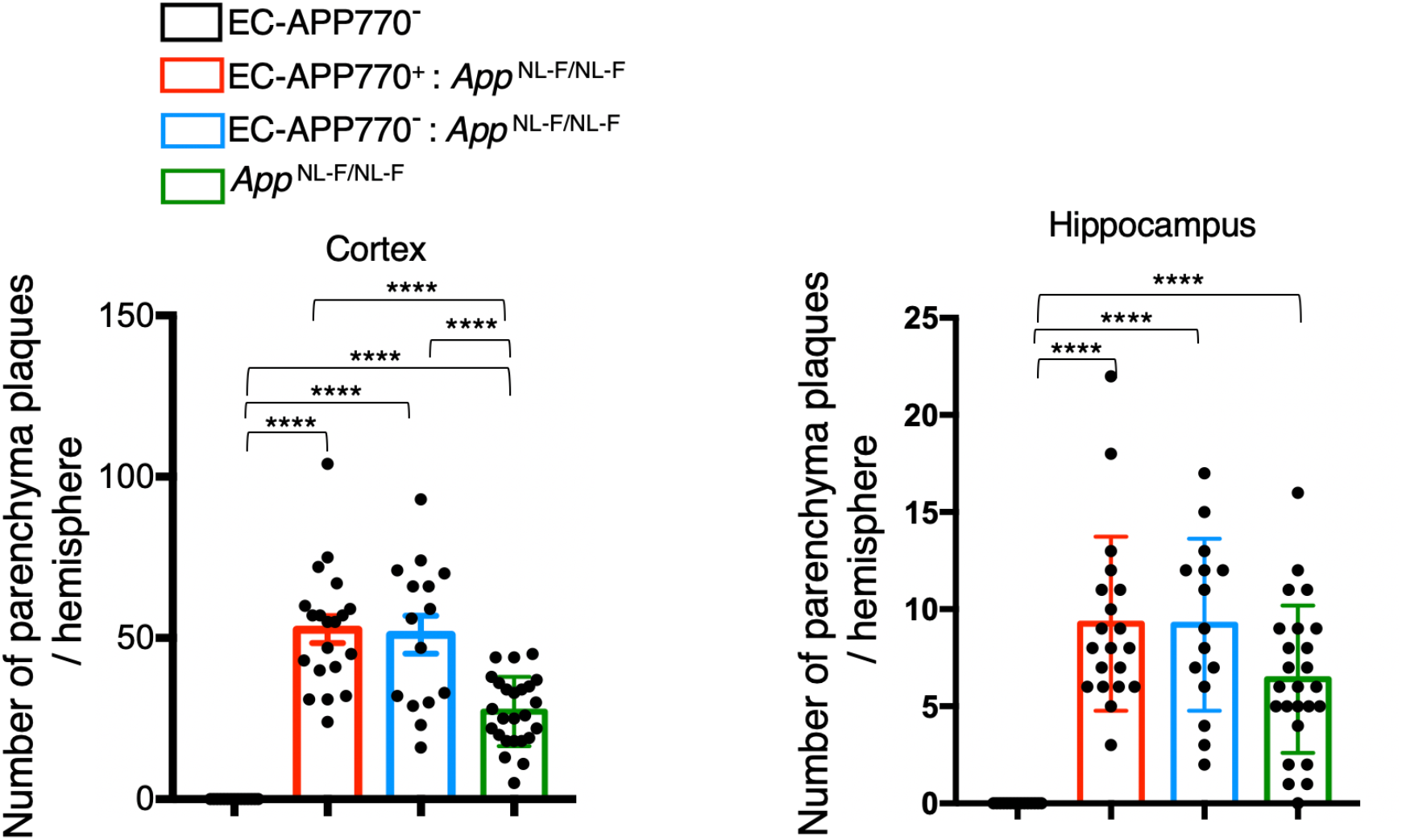
Similar parenchymal amyloid plaque deposition between *APP*^NL-F/NL-F^, EC-APP770^−^ and *APP*^NL-F/NL-F^ mice. The number of FSB-positive amyloid plaques in brain parenchyma of 15-month-old EC-APP770^+^, EC-APP770^+^ : *APP*^NL-F/NL-F^, EC-APP770^−^ : *APP*^NL-F/NL-F^ and *APP*^NL-F/NL-F^ mouse brain sections were quantified and shown as the mean±s.e.m. (n=4 mice; 5 sections were counted per mouse).

**Supplementary Figure 3.**
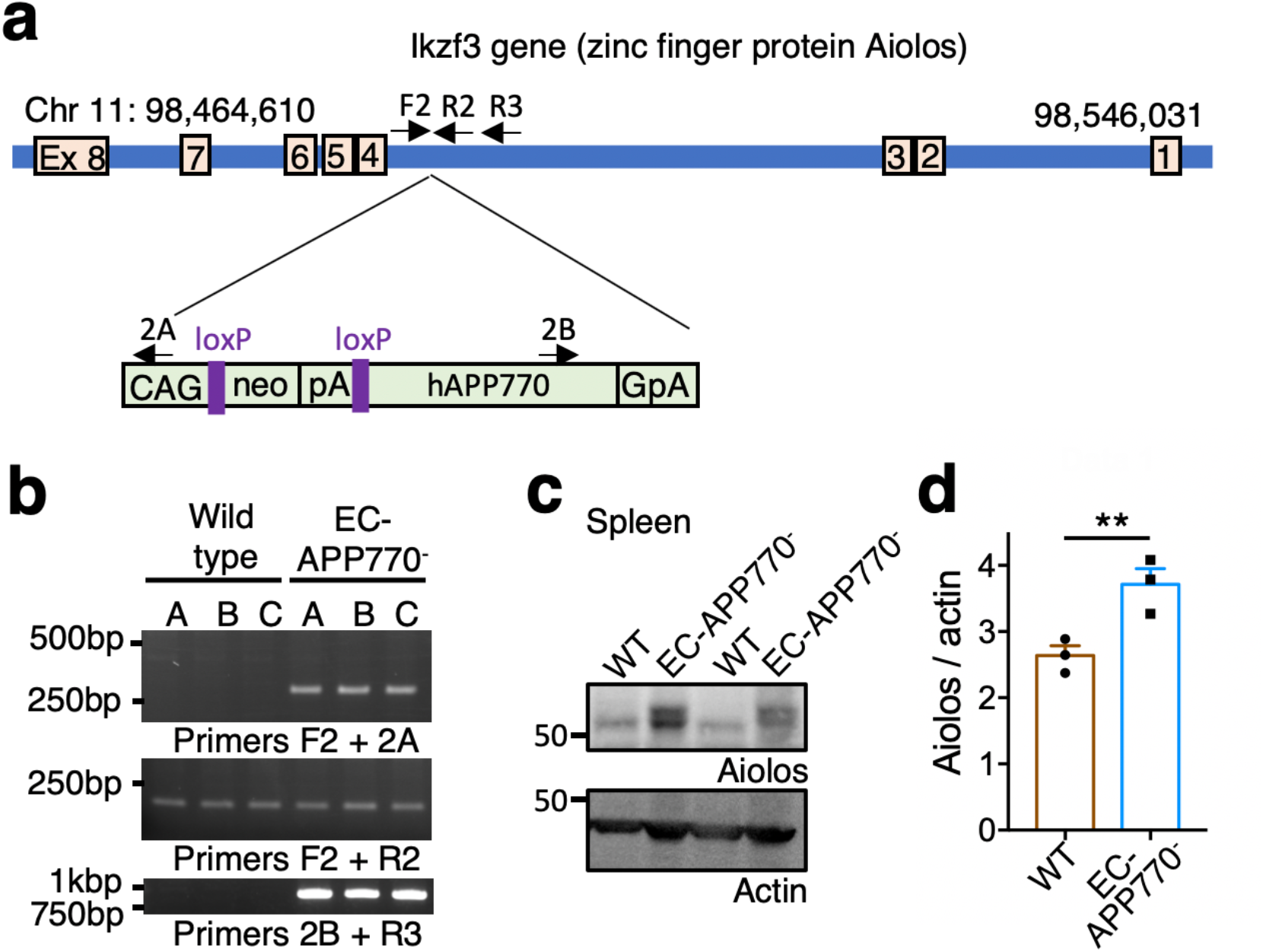
Identification of integration locus of hAPP770 transgene. (a) Schematic diagram showing hAPP770 transgene located in the Ikzf3 locus on chromosome 11. Positions of primers to validate the insertion site are also shown. (b) Insertion site was verified by PCR for left junction, inserted genome region, and right junction. (c) Spleen lysates from wild type (WT) and EC-APP^−^ mice were analyzed by western blot for Aiolos and actin. (d) The level of Aiolos in the spleens from both types of mice are shown as the mean±s.e.m. (n=3). Student’s *t*-test, p<0.01.

**Supplementary Figure 4.**
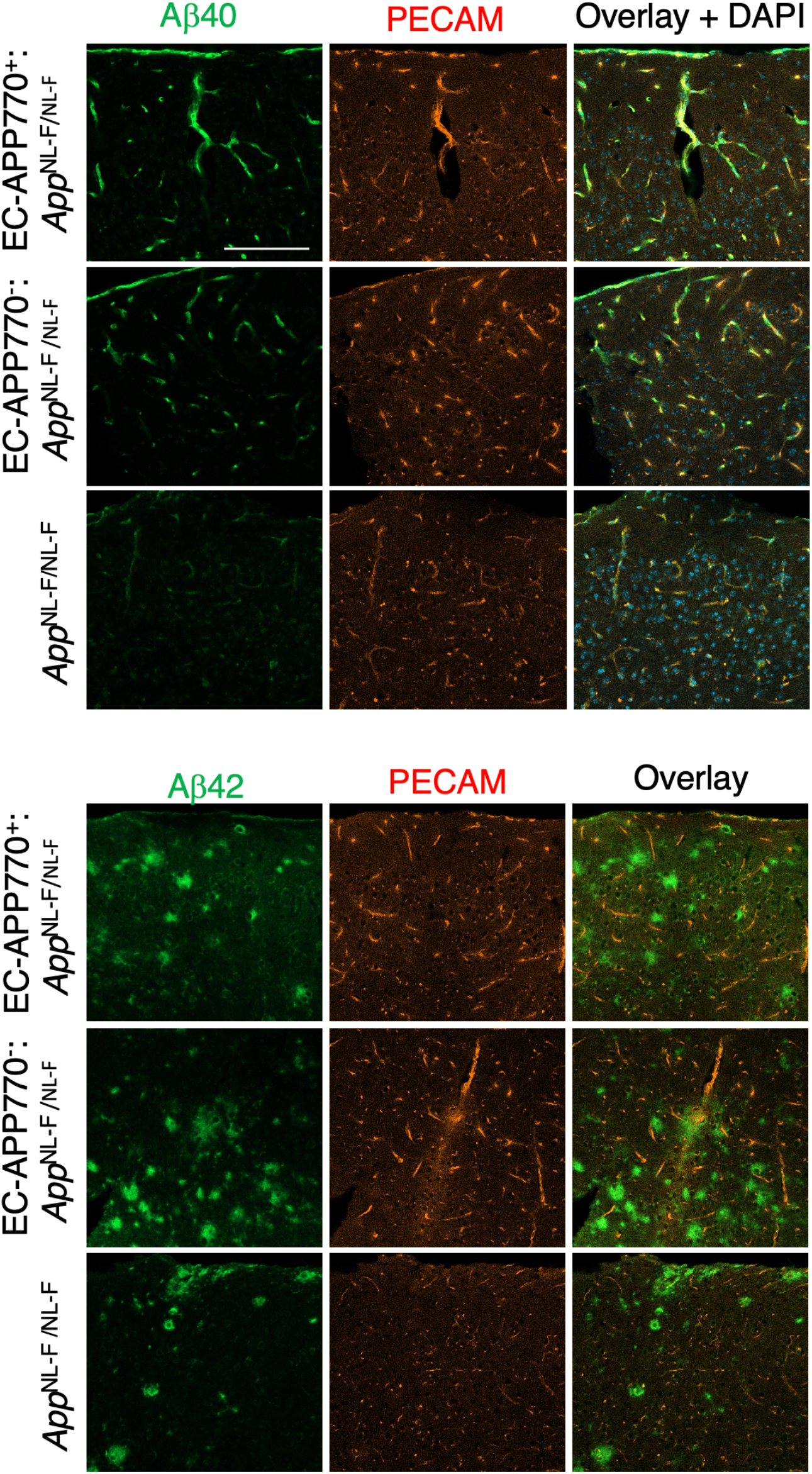
Aβ40 and Aβ42 are accumulated differently in the brains of EC-APP770^+^ : *APP*^NL-F/NL-F^ and *APP*^NL-F/NL-F^ mice. Brain cortical sections from 15-month-old EC-APP770^+^ : *APP*^NL-F/NL-F^, EC-APP770^−^ : *APP*^NL-F/NL-F^ and *APP*^NL-F/NL-F^ mice were stained for Aβ40 and Aβ42 with PECAM. Scale bar: 100 μm.

**Appendix Fig. 1d.**
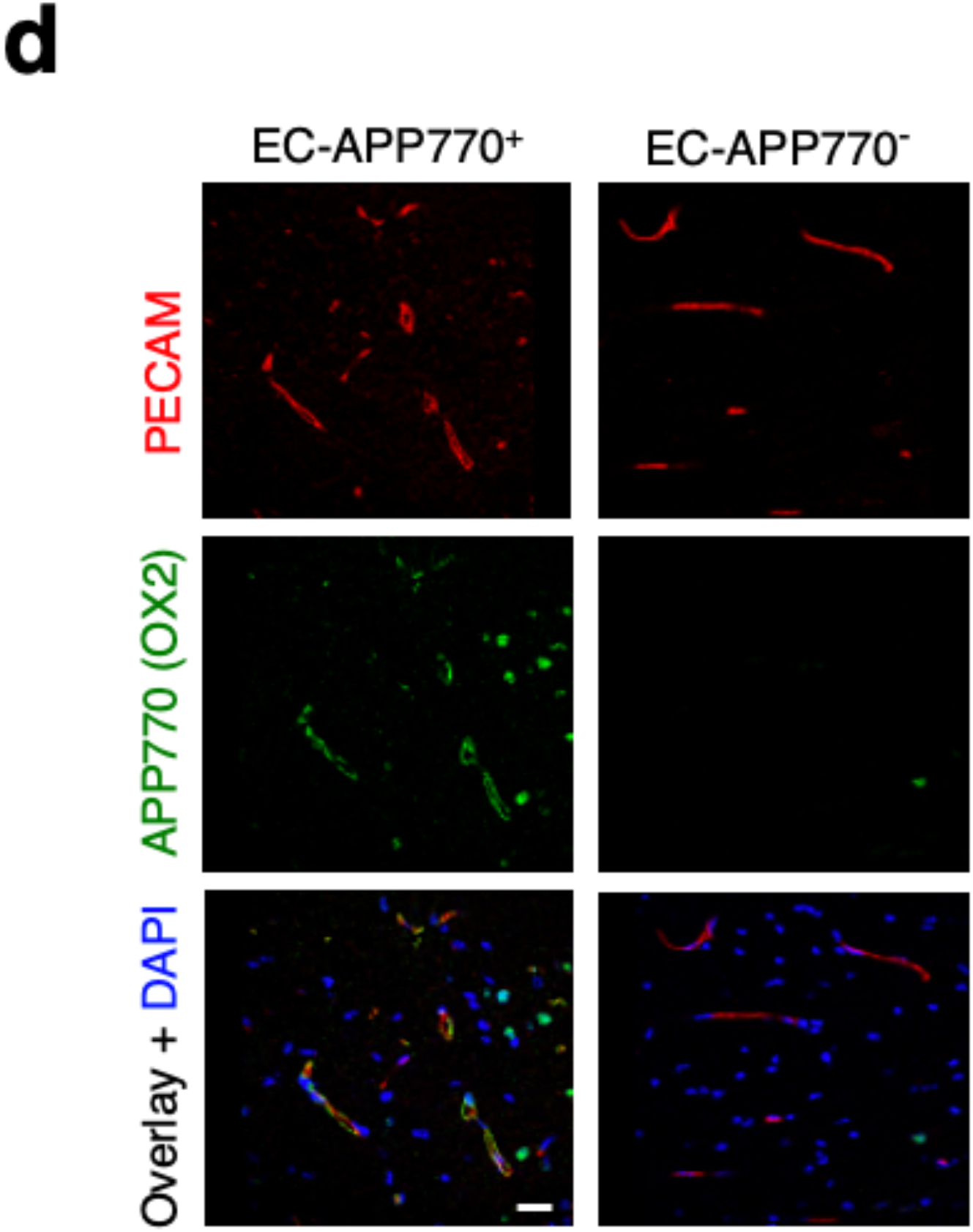

**Appendix Fig. 3d.**
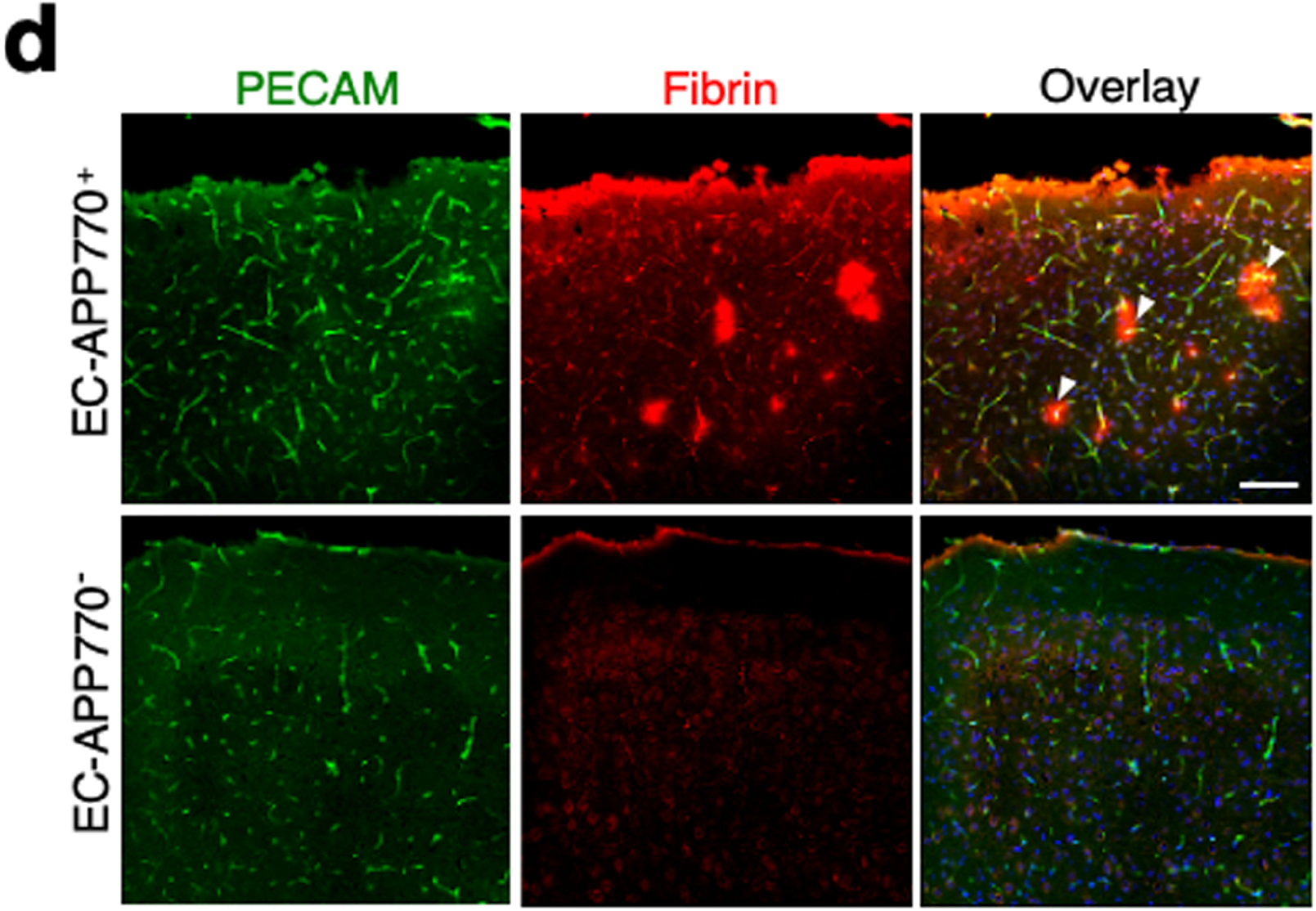

**Appendix Fig. 4a.**
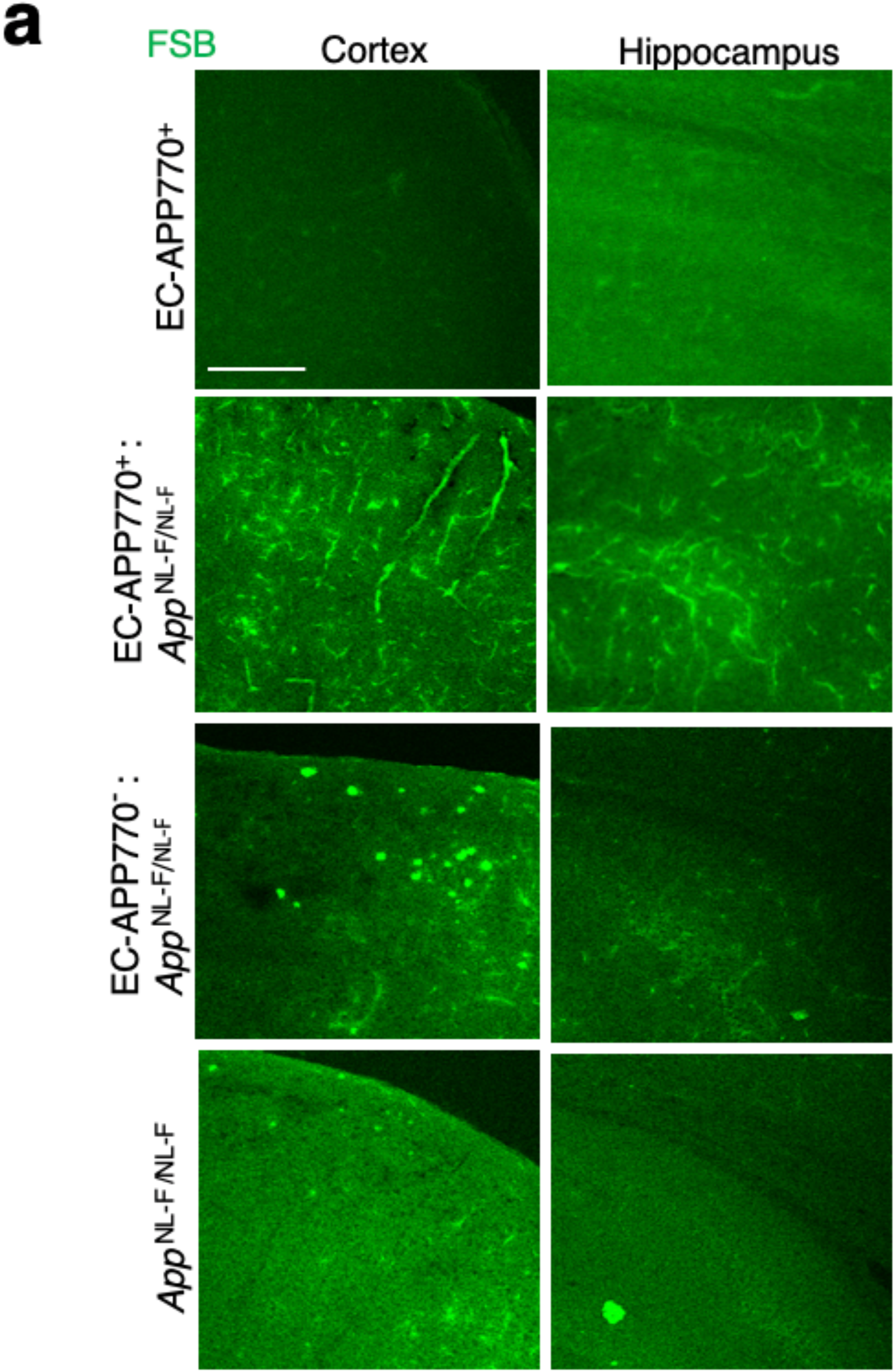

**Appendix Fig 4b, e.**
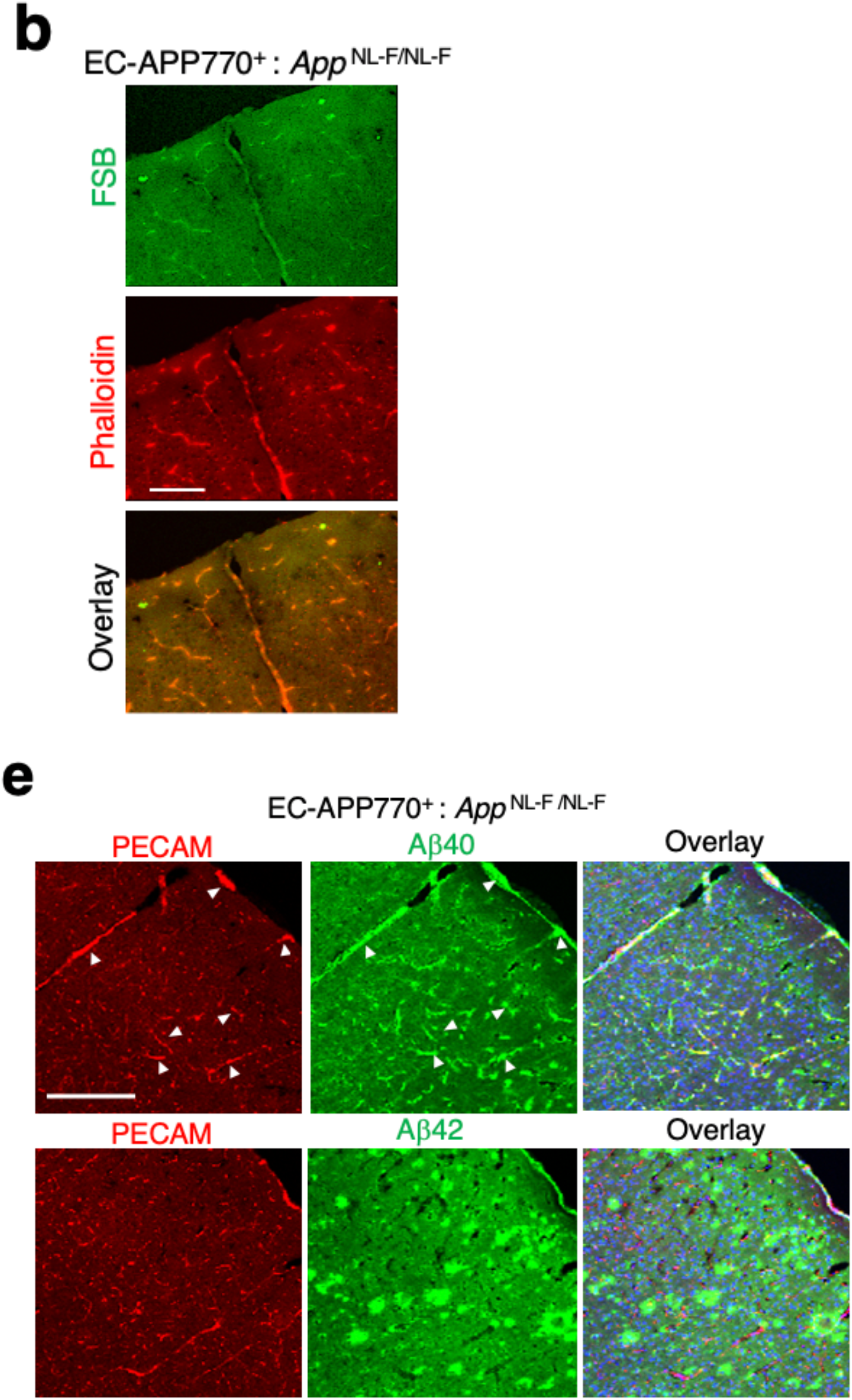

